# The Root of the Red Maple Paradox: Two distinct regeneration mechanisms determine leaf morphology, physiology, and competitive success in *Acer rubrum*

**DOI:** 10.1101/2022.03.01.482577

**Authors:** Laura Ostrowsky, Mark Ashton

## Abstract

Red maple is one of the most abundant and wide-ranging tree species in eastern North America and has rapidly increased in abundance over the past century following forest disturbances. This species is both a site- and light-generalist and has the unique ability to regenerate in forms that represent multiple stages of forest succession. However, red maple has modest physiological traits compared to its competitors, including low maximum photosynthetic rate, low photosynthetic nitrogen-use efficiency, and low foliar nutrient content. Red maple’s unremarkable physiology contradicts its competitive success. To untangle this paradox, we examine red maple’s two distinct regeneration mechanisms: seedlings and vegetative sprouts. Red maple can regenerate from seed, but can also sprout clonally from a stump following forest disturbance. We compare the morphology, physiology, and plasticity of these two regeneration mechanisms over 24 years of forest succession using a chronosequence of regenerating forest stands. We found that sprout-origin maples grow on average 6.5x taller and 5.5x faster than seed-origin maples. Sprout-origin trees display greater leaf spectral reflectance in the near-infrared range and greater stomatal density than seed-origin trees, demonstrating vegetative sprouts’ low water-use efficiency. Sprout-origin trees have more robust light-capture traits, including thicker palisade mesophyll than seed-origin trees. Sprouts also have a high plasticity between upper and lower leaves in many morphological and physiological traits, while seed-origin trees exhibited much less plasticity. The two distinct regeneration mechanisms of seed-origin and sprout-origin give rise to red maple trees with dramatically different leaf traits, allowing red maples to regenerate and thrive in a range of ecological conditions. Seed-origin maples are slow-growing, late successional, and shade-tolerant. Sprout-origin maples are fast-growing, early successional, and shade intolerant. This unique bimodal regeneration strategy may explain the red maple paradox and help predict forest composition and structure following disturbance.

## INTRODUCTION

Red maple (*Acer rubrum* L.) is one of the most far-ranging and ubiquitous tree species in eastern North America, and plays a unique ecological role in forest regeneration and succession as both a long-lived pioneer and a shade tolerant generalist (Fowells 1965; Burns and Honkala 1990). Its range and abundance have increased dramatically over the 20^th^ century, and it has increased in density as much as 6-fold in certain regions (Hardin et al. 2000; McDonald et al. 2002). Many second growth forests that have historically been dominated by oak (*Quercus* spp.) are shifting toward red maple, especially after partial disturbances or selective timber harvests that have removed the larger oak canopies (Nowacki et al. 1990; Smith and Vankat 1991; McWilliams et al. 2004; Fei et al. 2005). As a result, red maple is predicted to continue increasing in abundance in the future (Baah-Acheamfour et al. 2017; Sipe and Bazzaz 1995; Fei et al. 2005; Sanders-DeMott et al. 2017; Wheeler et al. 2017). Many studies have shown that red maple regenerates very effectively after disturbance and thrives in the face of biotic and abiotic factors associated with climate change, including herbivory, soil warming, reduced snowpack, increased freeze-thaw cycles (Sanders-DeMott et al. 2017), and increased disturbances causing large canopy gaps (Sipe and Bazzaz 1995; Fei et al. 2005; Wheeler et al. 2017). While relatives like sugar maple (*Acer saccharum* Marsh.) are projected to decline due to stresses related to climate change, red maple will likely increase in range and abundance (Sanders-DeMott et al. 2017).

Red maple’s dramatic increase and success has been puzzling foresters, ecologists, and plant physiologists for decades. Abrams (1998) referred to this puzzle as “the red maple paradox” in his review of the same name. He argues that this species presents a paradox because its mediocre physiological and morphological traits are at odds with its competitive success. Compared to competing tree species, red maple has low photosynthetic nitrogen use efficiency, low maximum and net photosynthetic rates, low foliar nitrogen concentration, low leaf mass per area, and unremarkable foliar structural characteristics (Abrams 1998, Nagel et al. 2002). In spite of these physiological traits, red maple increases branch numbers, leaf numbers, and total leaf area in gaps in comparison to other maple species (Sipe and Bazzaz 1994) and has much greater total biomass than other temperate species when grown competitively (Catovsky et al. 2002). Red maple has prolific seed production (Burns and Honkala 1990) and high levels of morphological plasticity (Sipe and Bazzaz 1994) compared to competing species. Red maple is also site generalist and light opportunist (Walters and Yawney 1990; Barnes and Wagner 2004). The ability to survive and compete in many edaphic conditions, light regimes, and stages of forest succession undoubtedly contributes to red maple’s prolific success. However, the physiological and morphological mechanisms that give rise to this generalist autecology are still unclear.

Given this physiological paradox, it is necessary to examine other traits within the species that may explain its success. One unusual feature of red maple is its ability to coppice, or sprout vegetatively from a stump. Following a forest disturbance that results in fallen or cut red maples, the remaining stumps will send up multiple sprouts in a multi-trunked growth form connected to the existing parent root system (hereafter referred to as “sprout-origin” red maples). Other red maples grow from seed and can persist as seedlings in the forest understory for years. Red maple therefore has two distinct regeneration mechanisms: seed-origin and sprout-origin trees. Roberts and Richardson (1985) proposed that these two recruitment mechanisms allow red maple to grow and succeed on a wide range of habitats. Many other researchers have drawn similar conclusions, and studies have shown that red maple can be represented at many stages of forest succession by originating from vegetative sprouts immediately after a disturbance or establishing in the understory through seed in later years (Ross et al. 1982; Roberts and Richardson 1985; Arthur et al. 1997; Tremblay et al. 2002). However, to our knowledge, no studies have examined and compared the physiological or morphological traits of these sprout-origin red maples to those maples of seed-origin. Studies of red maple physiology focus almost exclusively on seed-origin trees, leaving a gap in our understanding of red maple ecology. Very little is known about the functional differences between the seeding and sprouting regeneration mechanisms and their effects on forest regeneration. Additionally, no studies to date have examined red maple regeneration at temporal scales that allow for comparisons over early successional time – a period of critical competition and differentiation among species.

Our forests are changing rapidly in the face of climate change and increased human disturbance, and it is important to understand and anticipate forest composition shifts, species vulnerability, and regeneration patterns. Red maple plays a disproportionately large role in the regeneration of eastern forests and is unusually adept at responding to disturbance and stressors associated with climate change. Further research on red maple regeneration mechanisms is needed to understand the paradoxical strengths that allow this species to thrive and help foresters and ecologists anticipate the nature of forest regeneration after disturbance and over successional time.

In this study, we compare physiological and morphological traits of seed-origin and sprout-origin red maples over the first 24 years of forest succession. We hypothesize that seed-origin and sprout-origin red maples have distinct physiological and morphological traits, and that sprout-origin maples have a more robust and inefficient leaf physiology than seed-origin maples based on our observations of sprouts’ rapid early growth and their access to mature root systems.

## MATERIALS AND METHODS

### Study site

This research was conducted at Yale Myers Forest (YMF) (41.9529 N, 72.1239 W) in northeast Connecticut in May-June of 2018. YMF is a 3213 ha central hardwood-hemlock-pine forest (Westveld 1956). The elevation ranges between 170m and 300m above sea level and has a ridge-valley topography (Ashton and Larson 1996). The forest is second-growth, with dominant species including *Quercus rubra* L, *Pinus strobus* L, *Tsuga canadensis* L. (Carriere), *Acer spp*., *Betula spp*., and *Carya spp*. (Meyer and Plusnin 1945).

Two-thirds of YMF is managed for timber production and has been harvested as irregular shelterwoods over the past 30 years. A shelterwood harvest is a silvicultural technique that involves removing overstory trees in a stand but leaving reserve trees to purposely regenerate an even-aged cohort of individuals. Each year at Yale Myers, one or two stands are harvested in this manner, creating a chronosequence of stands that represent different successional stages (Duguid et al. 2016). This study used ten 4-17 hectare stands within the chronosequence all on similar ablation glacial till soil types defined as Charlton-Chatfield. They range in time since disturbance from two years (harvested in 2016) to 24 years (harvested in 1994).

### Sampling design and leaf collection

A total of 32 seed-origin and 32 sprout-origin red maples were identified in each of the ten shelterwood stands (for a total of 640 trees) and GPS coordinates were taken for each tree. All individuals in each site were in representative areas that accurately characterized the successional stage of development of the stand (i.e. not along edges or skid trails). A random number generator was used to select eight seed-origin and eight sprout-origin red maples for each of the 10 stands, for a total of 160 trees for measurement out of the original 640. Canopy cover over each seedling or sprout was measured using a densiometer. Height, root collar diameter, and diameter at breast height (1.37m) were measured for the seedlings. For the multi-trunked sprout-origin trees, we measured height of the tallest stem, number of stems, DBH of the six largest stems, and diameter at root collar of the parent stump.

All leaves were collected in late June through mid-July 2018 in a random order with respect to stand age. Four leaves were collected from each of the 160 trees: two upper leaves (a pair from the same node) and two lower leaves (also a pair from the same node). The upper leaves were the highest fully expanded leaves and the lower leaves were the lowest fully expanded leaves on the plant. On multi-trunked sprouts, all leaves were collected from the stem with the largest DBH.

Leaves were cut at the base of the petiole and kept in a cooler on ice until laboratory processing. The leaves in each pair were sorted into Group 1 and Group 2 for further measurement, so that the four leaves from each tree included a Group 1 upper leaf, a Group 1 lower leaf, a Group 2 upper leaf, and a Group 2 lower leaf. Physiology measurements (spectral reflectance and stomatal conductance) were performed within two hours of collection and all other leaf measurements were performed within 48 hours of collection.

### Leaf physiology and morphology measurements

Spectral reflectance and stomatal conductance were measured for all Group 1 leaves. Spectral reflectance was measured with a UniSpec Spectral Analysis System and UniWin V1.9 software to generate a reflectance curve of all wavelengths between 400 and 1100 nm for each leaf. Stomatal conductance of each leaf was measured with a Decagon Leaf Porometer (Decagon Devices, Inc., Pullman WA). Midrib length, petiole length, and leaf thickness were also measured for all Group 1 leaves. Leaf thickness was measured six times per leaf (left apical, left mid, left basal, right apical, right mid, and right basal) with a micrometer. A flatbed scanner was used to take an image of all Group 1 leaves and WinFOLIA software (Regent Instruments Inc.) measured the leaf perimeter, blade area, number of teeth, size of teeth, and lobing of each leaf.

All microscopy measurements were taken with a Keyence VHX-600 Digital Microscope. Stomatal size and density, and thickness of epidermal layers was measured for all Group 1 leaves. Stomatal size and density was measured at 1000x magnification. The underside of each leaf was cleaned with toluene and the number and diameter of all stomata in a 100×100 micrometer square was measured at four places on each leaf (left apical, left basal, right apical, and right basal). Epidermal layers were measured at 500x magnification. A cross-section of each leaf was cut and mounted, and the width of the upper epidermis, palisade mesophyll, spongy mesophyll, and lower epidermis was measured at three sections along the cross section. All measurements were taken from a 3D image of each location to account for uneven depth of the cross section.

### Measurements of relative water content

Relative water content (RWC) of all Group 2 leaves was calculated using fresh weight, saturated weight, and dry weight.

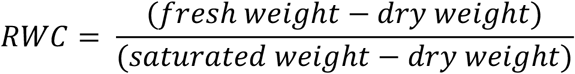

Fresh weight was measured immediately after leaf collection. For saturated weight, leaves were weighed after soaking in deionized water for 24 hours. The dry weight was measured after leaves dried for five days in a drying oven set to 70 degrees C.

### Statistical Analyses

All statistical analyses were performed using MiniTab18 and RStudio software (R Core Team 2018). To compare the four categorical groups (seedling upper, seedling lower, sprout upper, sprout lower), we fit a general linear model with stand age as a covariate and with interactions between tree type and leaf position, and between tree type and stand age (treating stand age as a categorical variable). We used one-way ANOVA tests with Tukey test for multiple comparisons to compare the means of each group.

Plasticity of morphological and physiological traits between upper and lower leaves was calculated using an index of phenotypic plasticity to compare mean values of traits across forest age classes and between regeneration mechanisms (adapted from Valladares et al. 2006).

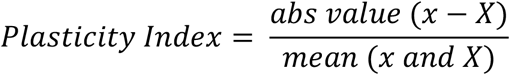

For each individual tree, a plasticity index was calculated for each trait between the upper and lower leaf from that tree. The plasticity indices for each trait were averaged across all trees tested to give an indication of variability in each trait between upper and lower leaves.

## RESULTS

Seed-origin and sprout-origin maples differed dramatically in morphology, physiology, growth, and plasticity across all variables tested (Table 1). Most variables also yielded significantly different results for upper versus lower leaves of each regeneration type. Sprout-origin trees were about 6.5x taller than seed-origin trees, and sprout-origin trees grew about 5.5x faster than seed-origin trees over the first 24 years of regeneration shown by the chronosequence of timber harvests.

**Table 1.**
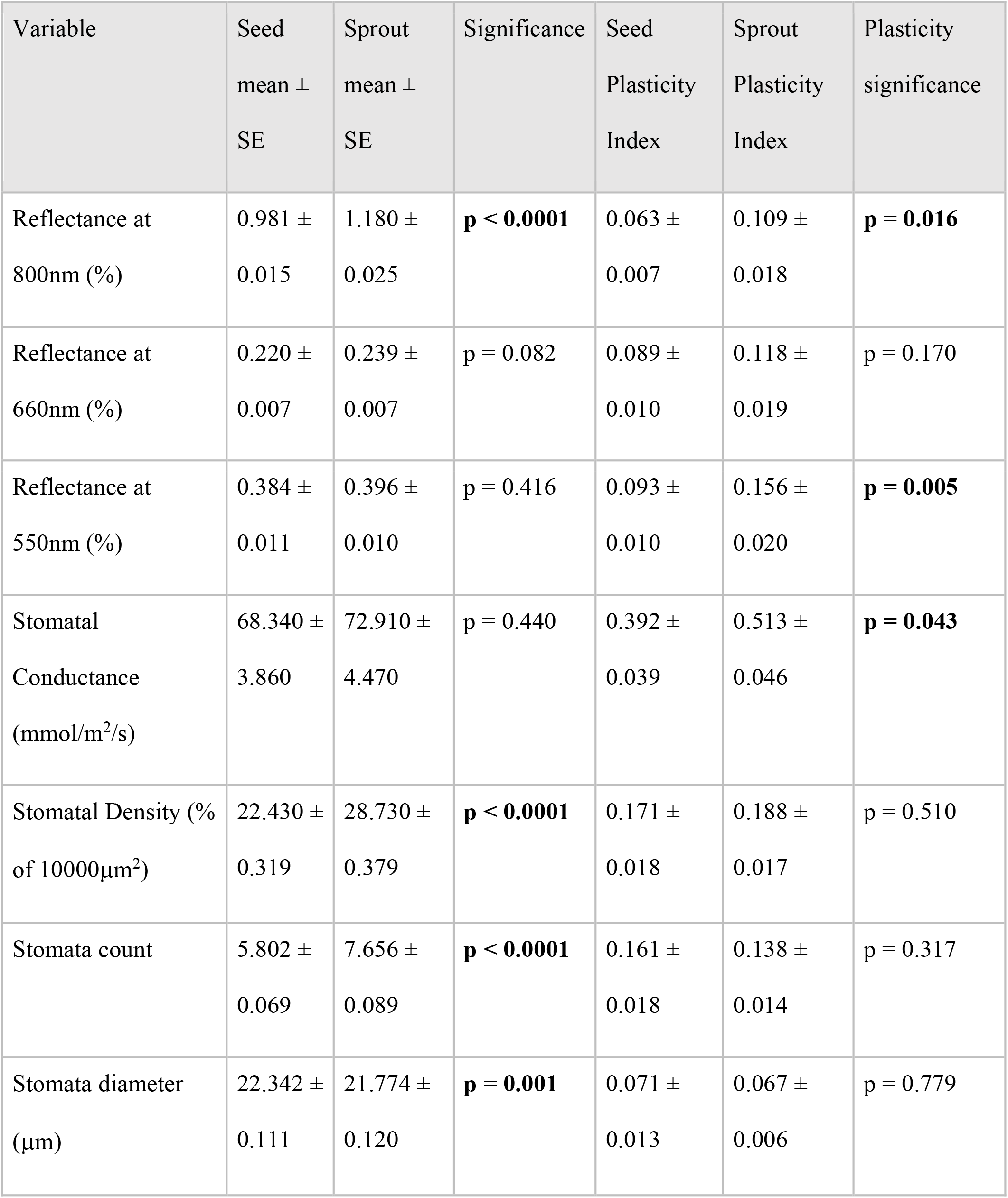
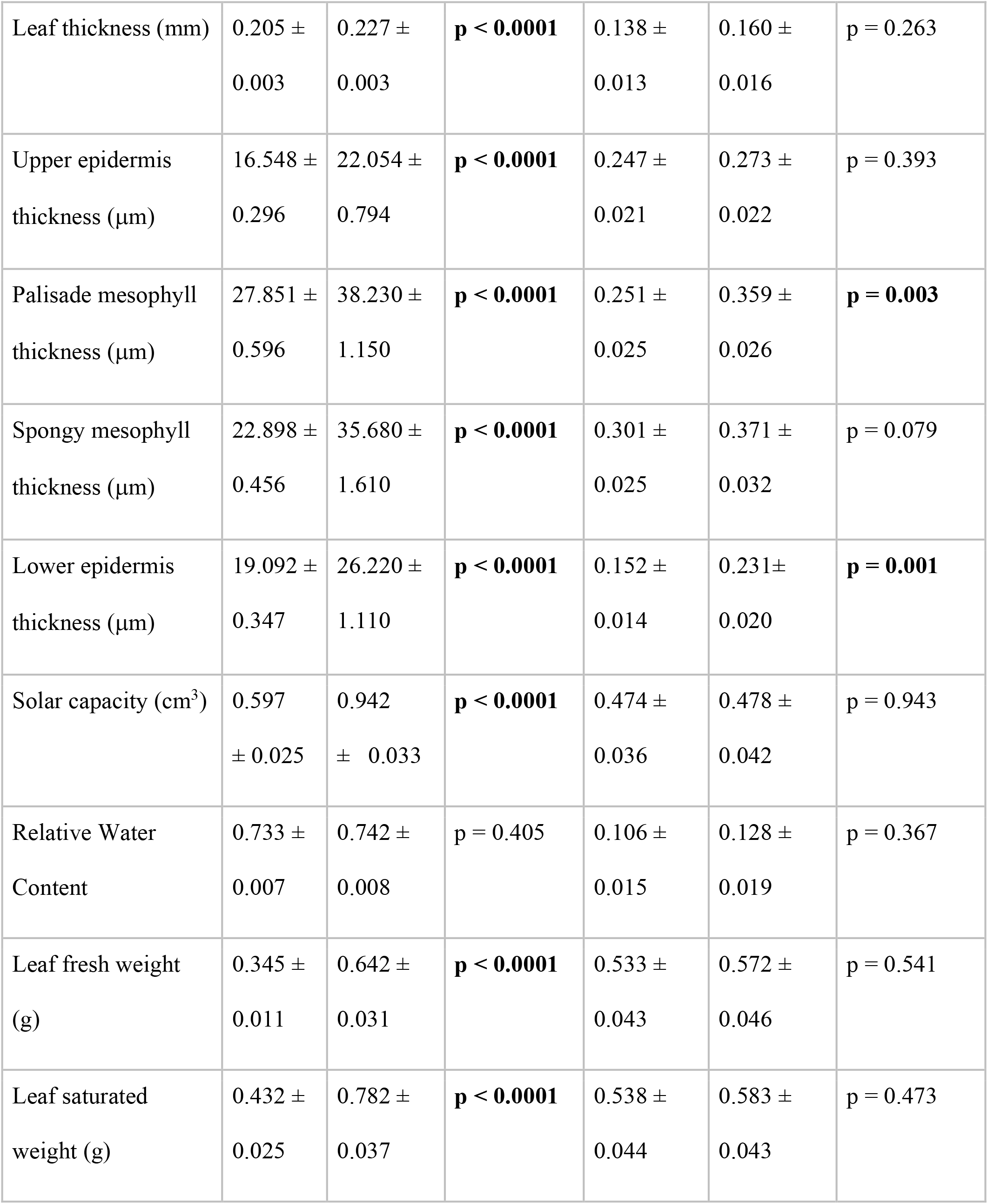
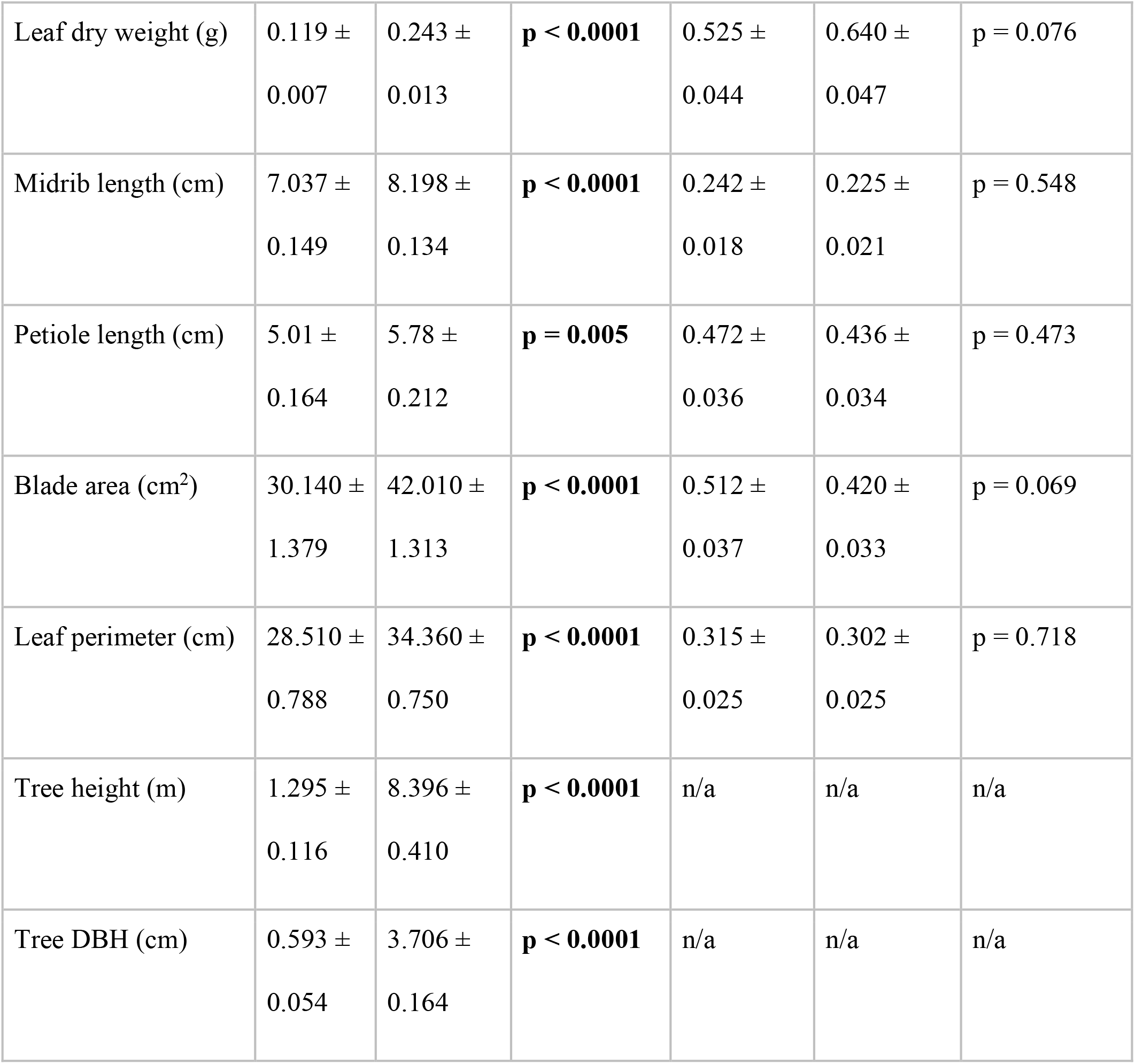
Summary of mean ± standard error of the mean for all primary variables tested (n = 320). Bold p-values represent statistically significant differences between the means of the seed and sprout groups.

### Leaf physiology and morphology measurements

Leaf spectral reflectance was greater in sprout-origin leaves than in seed-origin leaves at the wavelength 800nm (a representative wavelength for the far-red portion of the spectrum). There was no difference in reflectance between regeneration types for wavelengths 660 or 550. Stomatal density was higher in sprout-origin trees than in seedlings, and higher for upper leaves than lower leaves within sprout origin trees. The average diameter of stomata did not vary between groups, but the number of stomata per unit area was higher in sprout origin leaves than in seedling leaves.

Total leaf thickness varied by regeneration type (sprout vs. seedling) and leaf position (upper vs. lower). Sprout-origin trees had thicker leaves than seed-origin trees, and sprout-origin upper leaves were thicker than sprout-origin lower leaves. Total leaf thickness also had a negative relationship with stand age, as did each individual epidermal layer. Upper epidermis, palisade mesophyll, spongy mesophyll, and lower epidermis thickness were thicker in sprout-origin leaves than in seed-origin leaves, and thicker in upper leaves than in lower leaves within the sprout-origin leaves. Sprout-origin leaves also had longer leaf midribs than seedling leaves and longer petioles than seedling leaves. Sprout-origin leaves had significantly greater solar capacity (leaf thickness * blade area) than seed-origin leaves across all age classes, but solar capacity of sprout-origin leaves declined significantly over time with increasing shelterwood age at a rate of 1.8 x 10^−2^ cm^3^/year.

### Relative water content

Relative water content did not vary by regeneration type (seed-origin vs. sprout-origin). However, fresh weight, saturated weight, and dry weight were all higher for sprout-origin leaves than for seed-origin leaves. Fresh weight, saturated weight, and dry weight all also had a negative relationship to stand age (Multiple regression; R^2^ = 0.23).

### Plasticity

Sprout-origin leaves had a higher plasticity index than seed-origin leaves in five physiological and morphological leaf traits (Table 1). No other traits tested varied significantly in plasticity between seed-origin and sprout-origin leaves. The plasticity of spectral reflectance at 800nm was almost twice as high in sprout-origin leaves as it was in seed-origin leaves, and the palisade mesophyll thickness was about 1.5 times more plastic in sprout-origin than seed-origin leaves.

## DISCUSSION

Our results demonstrate that seed-origin and sprout-origin red maples are dramatically different in their leaf physiology, morphology, and plasticity. Across the 24 years of forest succession represented by the chronosequence, sprout-origin trees have lower water use efficiency, greater solar capacity, greater plasticity between upper and lower leaves, and much greater growth rates than seed-origin trees. These striking differences may explain the differences in regeneration time between sprout-origin trees and seed-origin trees and provide insight into studies that demonstrate very different roles that red maple can play within forest stand dynamics (Oliver and Stephens 1977; Oliver 1978; Palik and Pregitzer 1995; Tift and Fajvan 1999). We found that sprout-origin red maples dominate early in forest succession but diminish in vigor after 15 years of regeneration time, while seed-origin red maples remain in the understory and do not mature into mid-canopy trees until 15 or more years of regeneration. Together, these complementary strategies may explain red maple’s regenerative success and generalist abilities.

### Forest regeneration

Red maple is by far the most common tree species in Connecticut, and makes up 21% of the volume and 25% of the gross number of trees in the state (Butler 2017). It spreads particularly well following disturbance and land abandonment (Abrams 1998) and will likely continue to spread through forests of the northeast in the coming years. Red maple often dominates in the understory and dominates hardwood stands following disturbance, persisting 10-15 years longer than competing species during regeneration (Tift and Fajvan 1999). Its ability to regenerate by both seeds and sprouting may allow for this success. The ability to sprout vegetatively following disturbance has been adaptive for many tree species in the northeast (Ross et al. 1982), but it has been particularly adaptive for red maple. The seed-origin and sprout-origin trees in this study have dramatically different growth rates and forms over regeneration time. Sprout-origin trees grow over five times faster than seed-origin trees and the diameter at breast height (DBH) of each stem in sprout-origin trees is over twice as large as seed-origin DBH on average. Sprout-origin trees dominate in early succession but may begin to dwindle as they approach 15-20 years after regeneration (Oliver and Stephens 1977; Oliver 1978). In our study, the number of stems in their multi-trunked form declines from around 30 to around 3 or 4 as the stems outcompete one another, and the solar capacity of sprout-origin leaves declines steadily with shelterwood age. Meanwhile, the seed-origin trees finally become prominent and vigorous by the time they reach 15-20 years of regeneration, solidifying their place in mid-to-late succession (Palik and Pregitzer 1995; Tift and Fajvan 1999). This combination of strategies may allow red maple to proliferate so successfully in Connecticut’s forests.

### Water use efficiency

The high stomatal density (e.g. stomatal area index) and high reflectance in the infrared spectrum indicate that sprout-origin maples are very inefficient with water use, while seed-origin trees show high water-use efficiency with lower stomatal densities and lower infrared reflectance. Stomatal density can be indicative of a plant’s ability to adapt to changing environments and plastically regulate gas exchange (Franks and Beerling 2009; Franks et al. 2009). Stomatal area index is a function of both the number of stomata per unit area and the size of those stomata (Ashton and Berlyn 1994). The greater stomatal area index seen in the sprout-origin maple leaves is due entirely to a greater number of stomata, and not to any change in the size of stomata. We found that seed-origin and sprout-origin leaves have no difference in stomatal size, but seed-origin leaves have significantly fewer stomata per unit area than do sprout-origin leaves.

Reflectance in the infrared and near-infrared (NIR) spectrum relates to water-use efficiency because light interacts with water in the NIR wavelengths (Lorenzen and Jensen 1988; Smith et al. 2004). Several studies have demonstrated that NIR reflectance is highly correlated with leaf water content and chlorophyll content (Tucker 1977; Aase and Siddoway 1980; Tucker et al. 1981). We demonstrate that sprout-origin leaves have significantly higher reflectance at 800nm (in the NIR spectrum) than seed-origin leaves across all age classes, indicating that sprout-origin trees may have higher water contents in their leaves. Additionally, we found that sprout-origin trees have significantly larger leaves across all age classes. Because absolute water content relates to leaf size and leaf mass, sprout-origin trees may have even higher water content irrespective of differences in relative water content (Gonzalez and Gonzalez-Vilar 2001). Relative water content does not vary significantly between seed-origin and sprout-origin trees despite large differences in fresh weight, saturated weight, and dry weight (the three measures used to calculate RWC). This may indicate red maple’s strong ability to adapt leaf traits to conserve water and maintain a consistent water balance (Teulat et al. 2008).

Overall, the suite of traits relating to water indicate that sprout-origin red maples have a low water-use efficiency, which makes sense in light of their structural advantages; Sprouts have a full, mature root system from the beginning of their regeneration, affording them much easier access to water than the seed-origin red maples. This may be a considerable advantage as root system extension regulates plant water stress (Pereira and Chaves 1993). Seed-origin trees may be much more water limited than sprout-origin trees, and many other seed-origin leaf traits observed in this study may demonstrate a response to water limitation.

### Plasticity between upper and lower leaves

In addition to its unique regeneration mechanisms, red maple leaves can be highly plastic depending on the environment, changing in morphology and physiology between populations in valleys versus on ridges (Abrams and Kubiske 1990). Red maple seedling leaves grown in full sun have thicker leaves, cuticles, and epidermal and palisade mesophyll cell layers than leaves grown in the shade, and also have higher stomatal densities (Ashton et al. 1999). In this study we demonstrate that sprout-origin red maples have significantly higher plasticity between upper and lower leaves than seed-origin trees in many of the traits relating to water-use efficiency. In particular, sprout-origin trees are much more plastic in spectral reflectance at 800nm and 550nm and in their stomatal conductance. Early successional shade-intolerant species have been shown to exhibit greater leaf plasticity than later successional shade tolerant species (Ashton and Berlyn 1992; Ashton and Berlyn 1994). In our study, seed-origin and sprout-origin red maples are acting in a similar way, with shade tolerant, late successional seedlings exhibiting lower plasticity in water use traits, while early successional vegetative sprouts exhibit much higher plasticity in these traits.

### Light capture traits

Red maple’s light capture traits are very modest compared to traits in competing tree species. Low maximum and net photosynthetic rates and low photosynthetic nitrogen-use efficiency have been observed in many studies (Abrams 1998; Nagel et al. 2002), but clearly do not limit red maple’s prolific success. Nagel et al. (2002) found that red maple leaves have low construction cost and maintenance cost compared to the leaves of competing species and argued that these inexpensive leaves may offer some competitive advantage to explain red maple’s success. While this helps to explain red maple’s paradoxical success, it must be noted that studies of light-capture traits in red maple have been limited to seed-origin studies often in controlled environments. Sprout-origin red maples may not possess the same limitations, and more research is needed to determine if maintenance costs and construction costs are as low in sprout-origin red maple leaves as they are in seed-origin leaves.

Sprout-origin red maples may be better adapted to high light conditions and better equipped to process greater amounts of sunlight than seed-origin red maples due to their higher solar capacity. We define solar capacity as leaf thickness multiplied by blade area. Sprout-origin leaves have a significantly higher solar capacity than seed-origin leaves, although sprout-origin solar capacity declines significantly over time with stand development age while seed-origin solar capacity remains constant over time. Sprout-origin leaves also have significantly thicker upper epidermal layers, palisade mesophyll, spongy mesophyll, and lower epidermal layers than seed-origin leaves. The palisade mesophyll in particular is much thicker in sprout-origin leaves, indicating a higher capacity to process direct sunlight (Taiz and Zeiger 2014). The thickness of these epidermal and mesophyll layers and total leaf thickness decline over time with stand age in both seed-origin and sprout-origin leaves, and is potentially a response to the closing canopy as stands reach stem exclusion (Oliver and Larson 1996). Sprout-origin leaves also exhibit much greater plasticity in the thickness of these layers than do seed-origin leaves, demonstrating yet again their remarkable ability to adapt upper and lower leaves to varying light and water conditions.

### Conclusions

Together, two distinct regeneration mechanisms allow red maple to occupy multiple ecological niches and succeed both early and late in forest development following disturbance. Sprout-origin red maples act as long-lived pioneers, dominating in early succession and exhibiting extremely high levels of morphological plasticity. Seed-origin red maples exhibit a much more conservative strategy, maintaining a shade tolerant, water use efficient, and low-plasticity form in the understory for 15 or 20 years before beginning to grow to mid-canopy height. These drastic differences place seed-origin and sprout-origin red maples on opposite ends of a trait continuum and may explain the regeneration questions posed by the red maple paradox.

Red maple is a model of successful regeneration following disturbance. In an uncertain future of climate change and increased human disturbance, it is crucial to be able to anticipate which species will regenerate successfully and which may be at risk. Tree species without the ability to root-sprout, with low plasticity, and with specific niche partitioning may not be able to regenerate as effectively as those more similar to red maple in these traits. Red maple’s unique bimodal regeneration strategy sheds light on the red maple paradox and may help ecologists anticipate future forest composition shifts following disturbance.

## ACKNOWLEDGEMENTS

We thank Edward C. Armbrecht Jr. Family Fund, Carpenter Sperry Fund, Kohlberg-Donohoe Research Fellowship for supporting the field research conducted by the lead author on this study. We also thank the following colleagues for advice on both conducting the field research, providing equipment and logistical support, and providing comments on various versions of the manuscript: Dr. Graeme Berlyn, Dr. Marlyse Duguid, Dr. Craig Brodersen, Dr. Danica Doroski, David Woodbury, Caroline Tasirin, Ki’ila Salas, Jessica Wikle, Nicholas Olson, Kitty Kan, Marc Boudreaux, Danielle Losos, Jack Schleifer, Ananth Miller-Murthy, Paige Johnson, Ben Gross, and Mary Elizabeth Benton.

---

**Figure 1.**
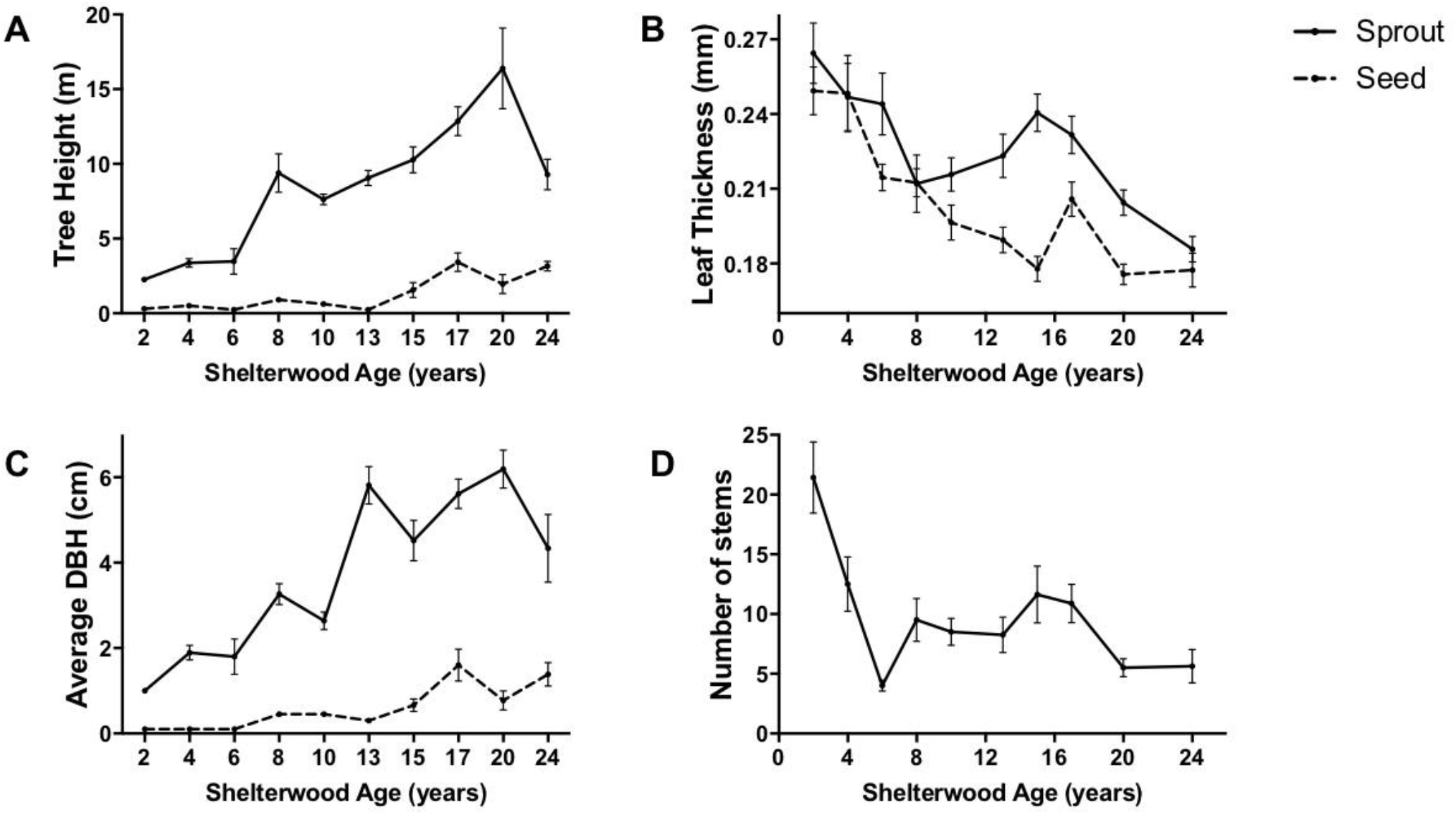
Mean tree height, leaf thickness, diameter at breast height (DBH), and number stems (in multi-trunked trees) over regeneration time in seed-origin and sprout-origin red maples (n = 160). Error bars represent standard error of the mean. Number of stems is measured for sprout-origin trees only, as seed-origin trees are not multi-stemmed.

**Figure 2.**
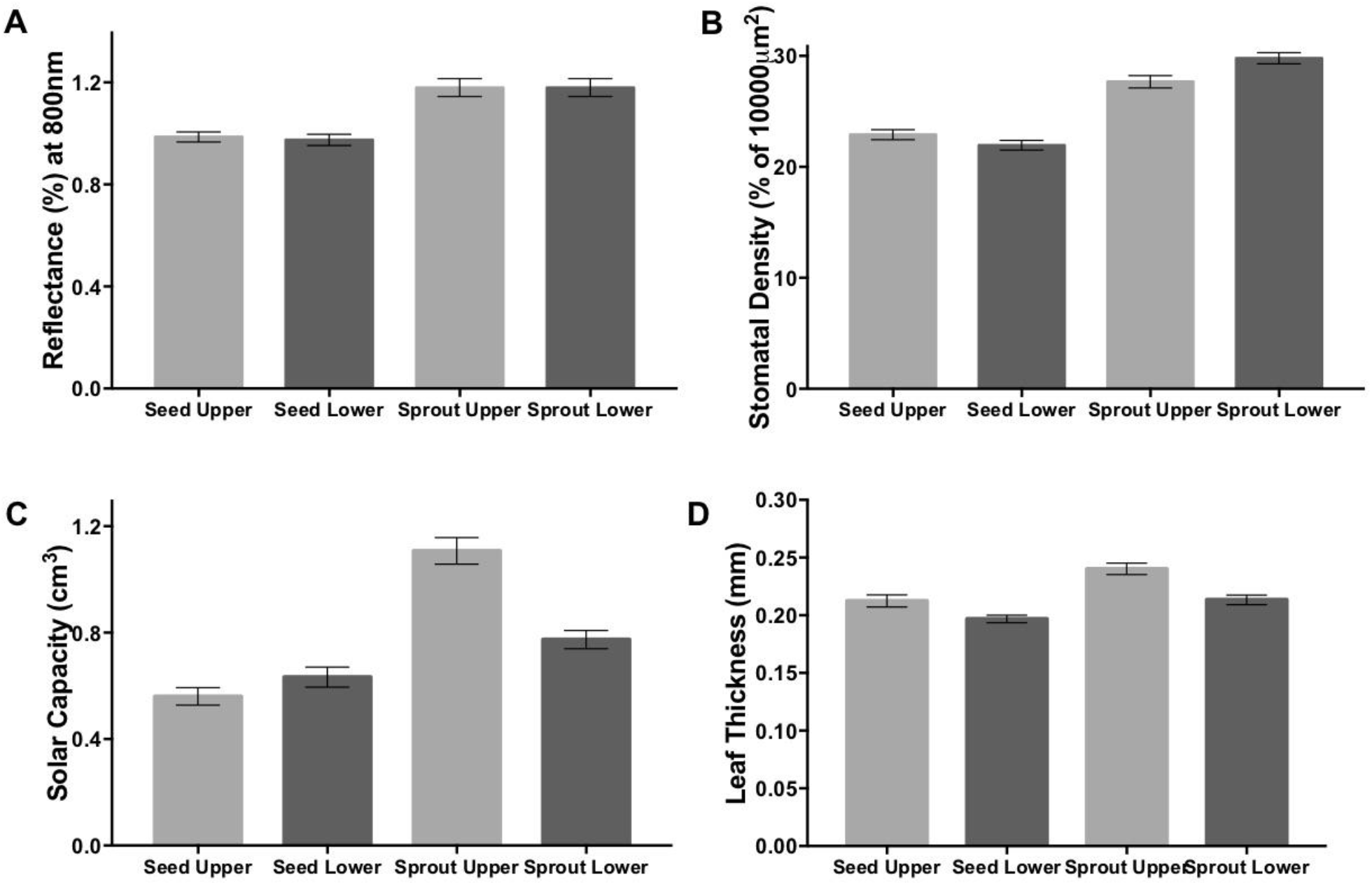
Comparisons of mean leaf traits between upper and lower leaves of seed-origin and sprout-origin red maples across all shelterwood ages (n = 320). Error bars represent standard error of the mean. Solar capacity is calculated as blade area * leaf thickness. Sprout-origin trees had significantly higher reflectance at 800nm, stomatal density, solar capacity, and leaf thickness than seed-origin trees.

**Figure 3.**
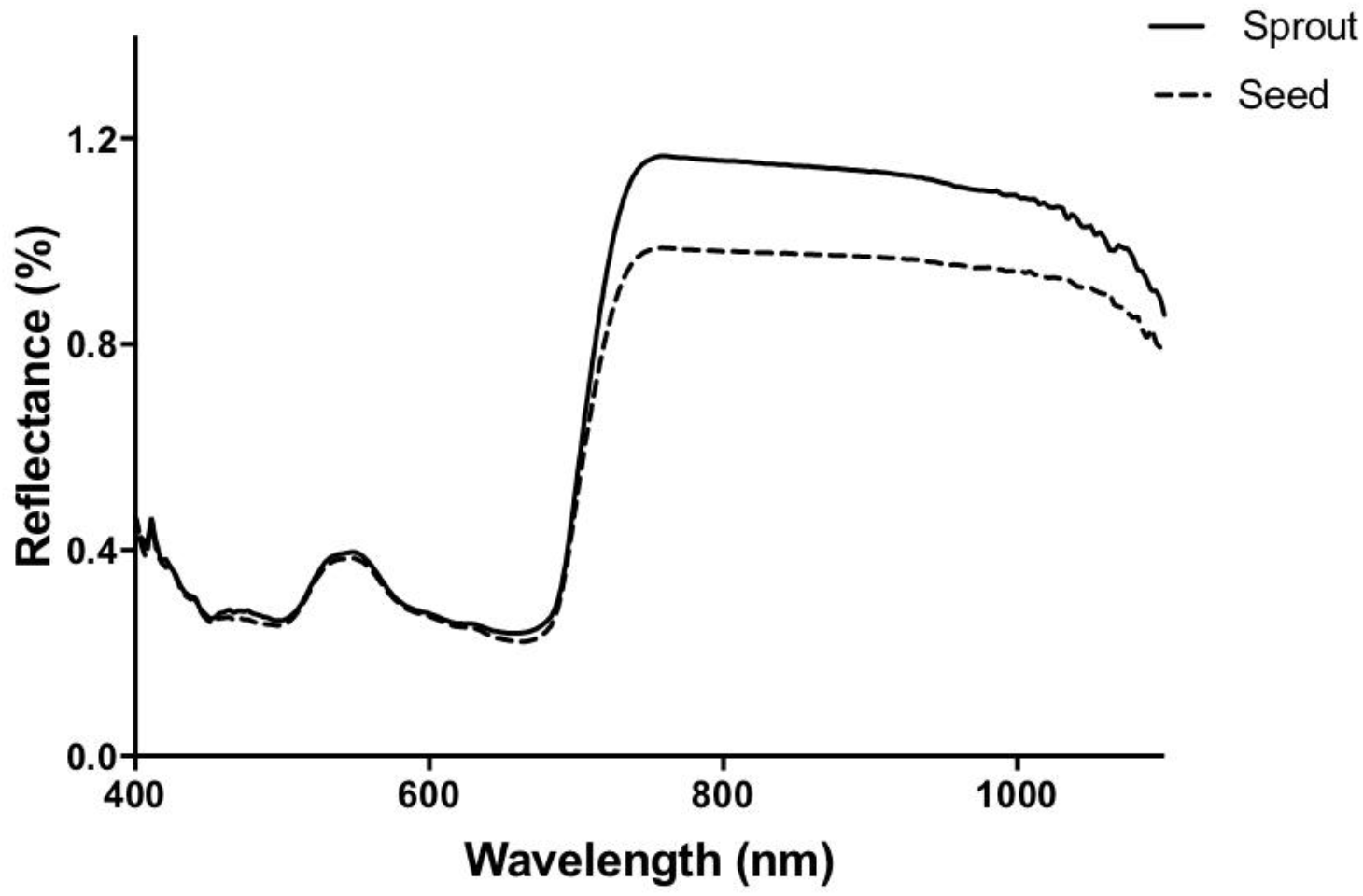
Mean spectral reflectance curves for all seed-origin and sprout-origin red maple leaves tested across all age classes (n = 320). Sprout-origin leaves had significantly higher spectral reflectance at 800nm wavelength than seed-origin leaves (GLM; p < 0.0001).

**Figure 4.**
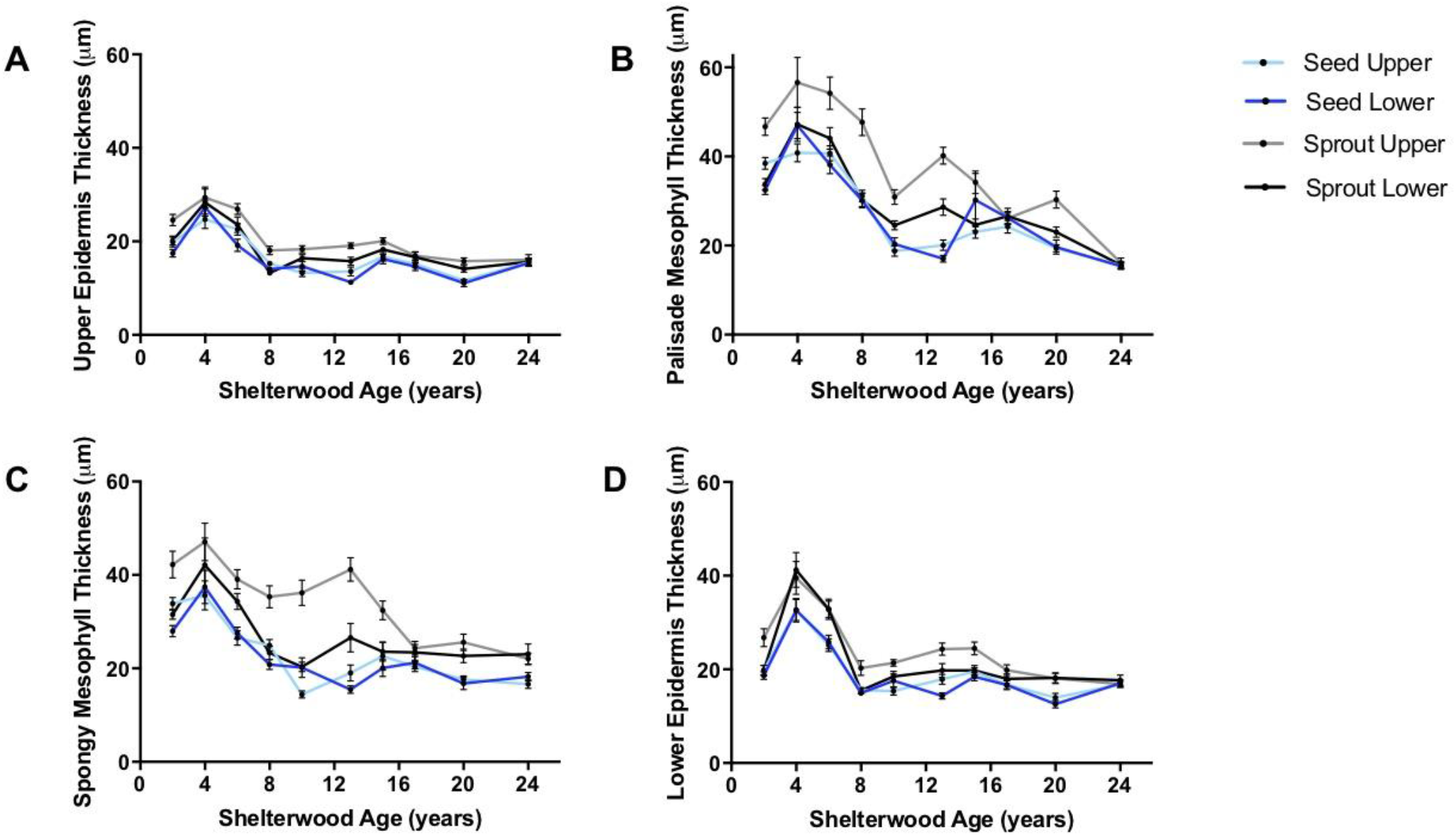
Mean thickness of leaf epidermal layers over regeneration time in upper and lower leaves of seed-origin and sprout-origin red maples (n = 960). Error bars represent standard error of the mean. Thickness of upper epidermis, palisade mesophyll, spongy mesophyll, and lower epidermis all decline with shelterwood age in both seed-origin and sprout-origin trees.

**Figure 5.**
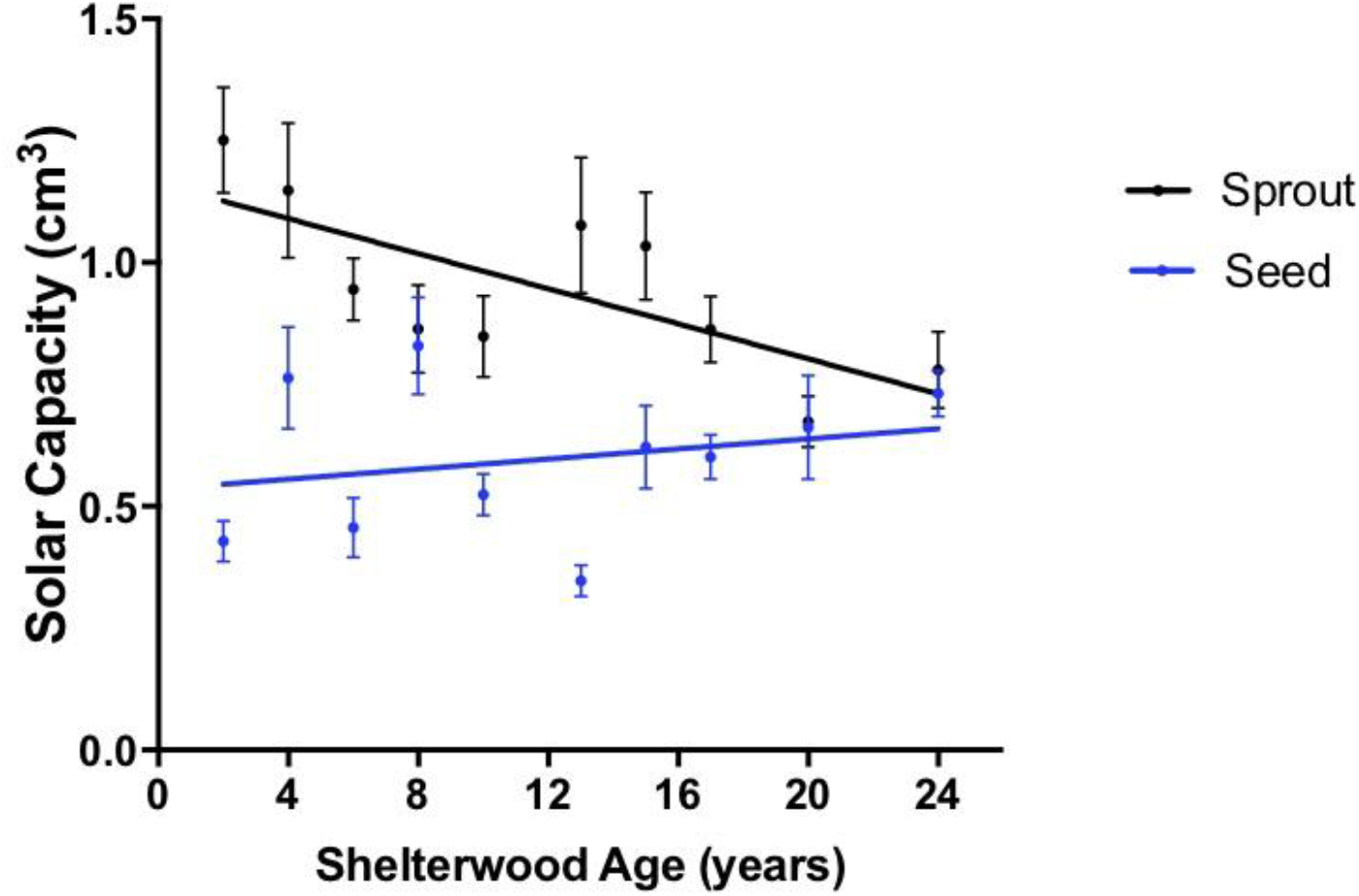
Solar capacity (blade area * leaf thickness) for seed-origin red maple leaves vs. sprout-origin red maple leaves over regeneration time (n = 320). Error bars represent standard error of the mean. Sprout-origin trees decrease in solar capacity at a rate of 0.018 cm3 per year. Slope for sprouts is significantly non-zero (p = 0.0173).

**Figure 6.**
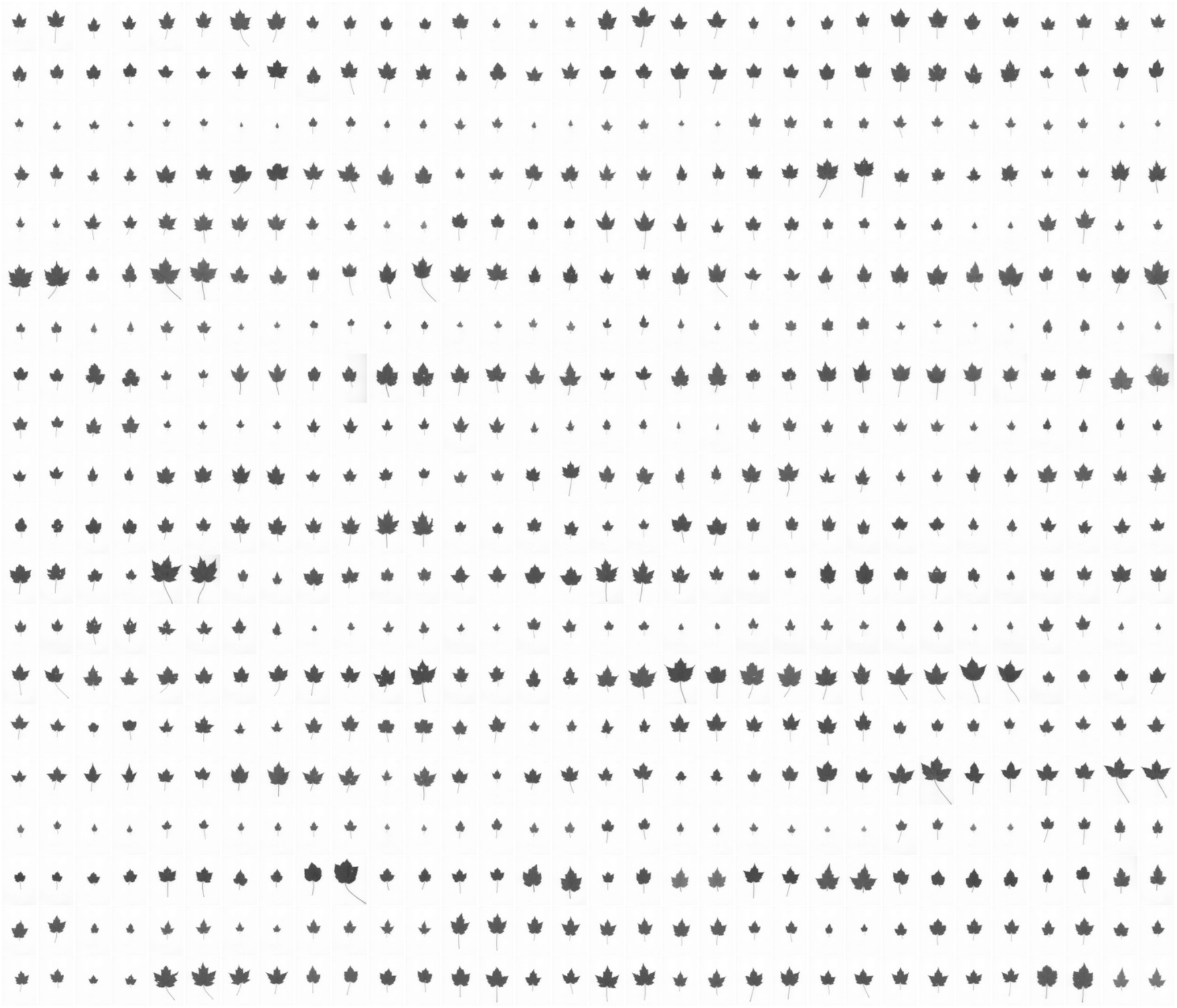
Compilation of 640 red maple leaf scans (to scale with each other) demonstrating the incredible variety in leaf morphology across samples. Upper and lower leaves from seed-origin and sprout-origin red maples across all ten shelterwood sites are represented.

